# Mercury and water level fluctuations in lakes of northern Minnesota

**DOI:** 10.1101/118638

**Authors:** James H. Larson, Ryan P. Maki, Victoria G. Christensen, Mark B. Sandheinrich, Jaime F. LeDuc, Claire Kissane, Brent C. Knights

## Abstract

Large lake ecosystems support a variety of ecosystem services in surrounding communities, including recreational and commercial fishing. However, many northern temperate fisheries are contaminated by mercury. Annual variation in mercury accumulation in fish has previously been linked to water level (WL) fluctuations, opening the possibility of regulating water levels in a manner that minimizes or reduces mercury contamination in fisheries. Here, we compiled a long-term dataset (1997–2015) of mercury content in young-of-year Yellow Perch (*Perca flavescens*) from six lakes on the border between the U.S. and Canada and examined whether mercury content appeared to be related to several metrics of WL fluctuation (e.g., spring WL rise, annual maximum WL, and year-to-year change in maximum WL). Using simple correlation analysis, several WL metrics appear to be strongly correlated to Yellow Perch mercury content, although the strength of these correlations varies by lake. We also used many WL metrics, water quality measurements, temperature and annual deposition data to build predictive models using partial least squared regression (PLSR) analysis for each lake. These PLSR models showed some variation among lakes, but also supported strong associations between WL fluctuations and annual variation in Yellow Perch mercury content. The study lakes underwent a modest change in WL management in 2000, when winter WL minimums were increased by about 1 m in five of the six study lakes. Using the PLSR models, we estimated how this change in WL management would have affected Yellow Perch mercury content. For four of the study lakes, the change in WL management that occurred in 2000 likely reduced Yellow Perch mercury content, relative to the previous WL management regime.

## Introduction

Large lake ecosystems support a variety of ecosystem services, including commercial harvest, sport fishing and tourism (Holmlund and Hammer 1999). However, many northern temperate fish communities are contaminated with methylmercury, which harm fish and may affect humans and other fish consumers (Scheuhammer et al. 2007). Reducing mercury contamination in fisheries is a major management goal of many natural resource managers (e.g., Voyageurs National Park).

The amount of mercury in the biosphere has increased globally as a result of human activities and atmospheric mercury can travel long distances from its source (Munthe et al.2007). Even otherwise pristine locations may receive substantial atmospheric deposition of mercury (Driscoll et al. 2007). A portion of this atmospherically deposited mercury (often in oxidized elemental form) may then be converted to methylmercury (CH3HG_+_) by sulfate reducing bacteria in anoxic sediments (Selin 2009). Methylmercury can then enter the food web and accumulate to much greater concentrations in higher trophic levels (Selin 2009).

Methylmercury production is often measured using a sentinel species approach. The sentinel species approach involves sampling a single species over space or time, usually at the same age or life stage. In previous studies, the young-of-year of a species that feeds at lower trophic levels has been used, so that the temporal variation in mercury concentrations can be tied to a particular season or exposure period (Sorensen et al. 2005, Wiener et al. 2006,Larson et al. 2014). Using these methods, controls over spatial variation in methylmercury production have been inferred, leading to the conclusion that methylmercury production is higher in watersheds with abundant wetlands (Wiener et al. 2006,Selin 2009). Fewer studies have identified controls over temporal variation in methylmercury production, but among those that have, water level (WL) fluctuations have been identified as a potentially important driver of annual and longer-term variation in methylmercury production (Sorensen et al. 2005, Dembkowski et al. 2014a, Larson et al. 2014,Willacker et al. 2016). Water levels in lakes and reservoirs are often managed for multiple uses, hence water level management could be a tool used to reduce methylmercury production (Mailman et al. 2006).

In an analysis of lakes in Minnesota, Sorensen et al. (2005)found strong associations between WL fluctuations and mercury concentrations in young-of-year Yellow Perch (*Perca flavescens*). A follow-up study by Larson et al. (2014)found that these associations varied in magnitude spatially, with some lakes having no WL-mercury associations and others having very strong associations. Here, our objective was to use multivariate methods to identify WL-mercury relationships on a lake-specific basis. To accomplish this objective, we analyzed annual measurements of mercury in tissues of young-of-year Yellow Perch from six large lakes during 1997–2015, some of which have been used in previous studies (Sorensen et al. 2005,Larson et al. 2014).

## Methods

### Study Sites

The Rainy-Namakan Reservoir complex occurs on the border between the United States and Canada. The complex consists of several naturally occurring lakes that have been impounded. Water levels are regulated for multiple purposes (e.g., hydropower, flood control, fisheries). Thirteen sites in six lakes from the Rainy-Namakan complex in or near Voyageurs National Park (Minnesota, USA) were sampled between 2013 and 2015 (Crane Lake, Lake Kabetogama, Little Vermilion Lake, Namakan Lake, Rainy Lake and Sand Point Lake; Table 1). Fish were collected between mid-September and early November each year. Fish were collected with 15.2 or 30.5 m bag seines with 6.4 mm mesh (bar), following procedures approved by the National Park Service’s Institutional Animal Care and Use Committee.

**Table 1.**
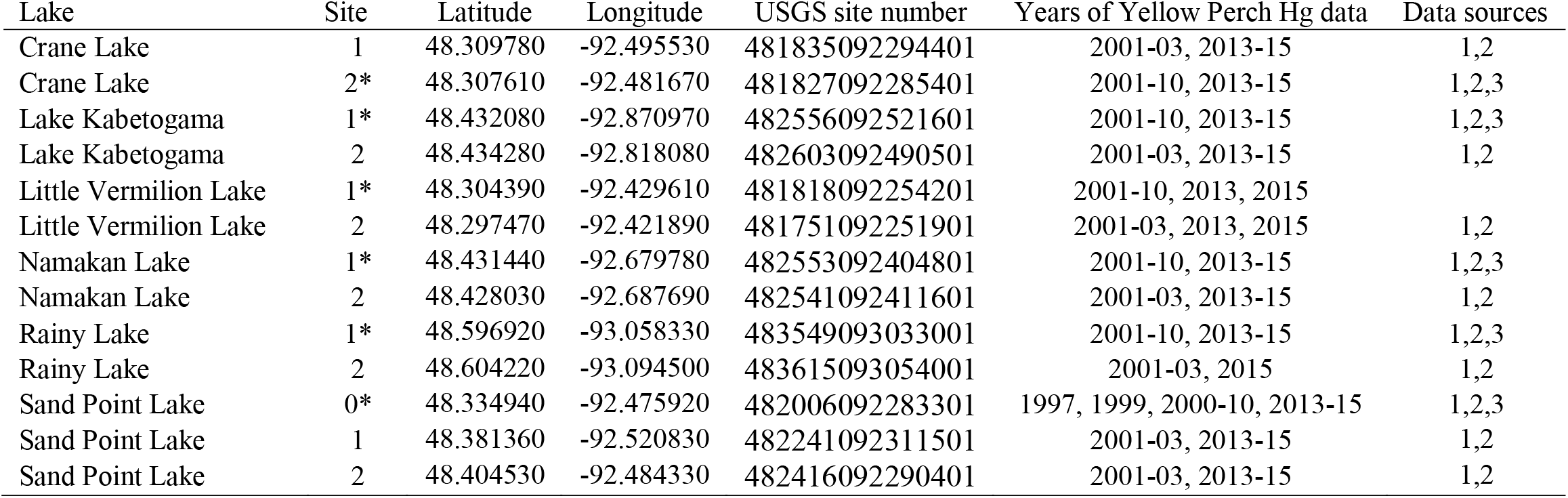
Sites from which young-of-year Yellow Perch were captured and mercury content measured. USGS site numbers refer to site data available through the website https://waterdata.usgs.gov/nwis. Data sources are 1-Sorensen et al. 2005, 2-Christensen et al. 2017 and 3-Larson et al. 2014.

| Lake | Site | Latitude | Longitude | USGS site number | Years of Yellow Perch Hg data | Data sources |
| --- | --- | --- | --- | --- | --- | --- |
| Crane Lake | 1 | 48.309780 | -92.495530 | 481835092294401 | 2001-03, 2013-15 | 1,2 |
| Crane Lake | 2* | 48.307610 | -92.481670 | 481827092285401 | 2001-10, 2013-15 | 1,2,3 |
| Lake Kabetogama | 1* | 48.432080 | -92.870970 | 482556092521601 | 2001-10, 2013-15 | 1,2,3 |
| Lake Kabetogama | 2 | 48.434280 | -92.818080 | 482603092490501 | 2001-03, 2013-15 | 1,2 |
| Little Vermilion Lake | 1* | 48.304390 | -92.429610 | 481818092254201 | 2001-10, 2013, 2015 |  |
| Little Vermilion Lake | 2 | 48.297470 | -92.421890 | 481751092251901 | 2001-03, 2013, 2015 | 1,2 |
| Namakan Lake | 1* | 48.431440 | -92.679780 | 482553092404801 | 2001-10, 2013-15 | 1,2,3 |
| Namakan Lake | 2 | 48.428030 | -92.687690 | 482541092411601 | 2001-03, 2013-15 | 1,2 |
| Rainy Lake | 1* | 48.596920 | -93.058330 | 483549093033001 | 2001-10, 2013-15 | 1,2,3 |
| Rainy Lake | 2 | 48.604220 | -93.094500 | 483615093054001 | 2001-03, 2015 | 1,2 |
| Sand Point Lake | 0* | 48.334940 | -92.475920 | 482006092283301 | 1997, 1999, 2000-10, 2013-15 | 1,2,3 |
| Sand Point Lake | 1 | 48.381360 | -92.520830 | 482241092311501 | 2001-03, 2013-15 | 1,2 |
| Sand Point Lake | 2 | 48.404530 | -92.484330 | 482416092290401 | 2001-03, 2013-15 | 1,2 |

### Water level estimates

Water level metrics calculated here were similar to those used in Sorensen et al. (2005)and Larson et al. (2014). All water level data were obtained from the Lake of the Woods Water Control Board, which provided daily averages for the entire study period. Within-year minimum WL and maximum WL were used to calculate water level rise (WLR). Mean and standard deviation (SD) in daily WL were also calculated. In addition, change in maximum water level (ΔmaxWL) from the previous year was calculated. Water levels in the Rainy-Namakan lake complex follow a seasonal pattern, with early-spring minimums and early to mid-summer maximums. The WLR, minimum, maximum, mean and SD in WL were calculated for the entire year, for the spring (April-June) and for the summer (July-September).

### Atmospheric deposition of mercury and sulfate

Mercury and sulfate deposition were measured by the National Atmospheric Deposition Program (NADP). For mercury, annual data from the Fernberg monitoring location (MN18) were obtained from the NADP website (NADP 2012a). For sulfate, annual data from the Sullivan Bay monitoring location (MN32) were also collected from the NADP website (NADP 2012b).

### Temperature Data

Water temperature data were not available for most of the lakes over the time scales needed for analysis. Instead, air temperatures and equation 1 from Chezik et al. (2014)were used to calculate annual degree days for each lake. All data were obtained from the National Oceanic and Atmospheric Administration's National Climatic Data Center (http://www.ncdc.noaa.gov/data-access). Station USW00014918 was used for all of the lakes in this study. In this formulation, the degree days for a single day (DD; °C-days) are calculated as:

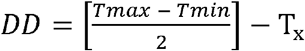

T_x_ is the threshold temperature under consideration (e.g., 0°C or 5°C) and T_max_ and T_min_ are the daily maximum and minimum temperatures respectively. Negative daily DD estimates are discarded, and the positive daily DD estimates are summed for the year. We calculated DDs for a T_x_ of 0°C, 5°C, 10°C and 15°C.

### Fish mercury analysis

Fish mercury data is compiled from three previously published studies:Sorensen et al. (2005), Larson et al. (2014)and Christensen et al. (2017,Table 1). Methods for estimating the mercury content for young-of-year Yellow Perch were reported in those manuscripts. For this analysis, we used only mercury content for young-of-year Yellow Perch on a dry mass basis.

### Statistical Methods

Data were compiled using a combination of R Version 3.1.0 (R Development Core Team 2014). All statistics were completed in R.

We used partial least squared regression (PLSR) to identify associations between young-of-year Yellow Perch mercury content and WL fluctuations. Partial least squared regression is similar to principal components analysis (PCA;Manly 2005), in that axes of co-variation among variables are identified. However, PLSR has a predictor-response structure that is used to identify the components (Carrascal et al. 2009). Partial least squared regression is useful in cases where sample sizes are low and many predictor variables that are strongly correlated are believed to be important (Carrascal et al. 2009). Essentially, PLSR identifies components of variation in the predictor variables (presumed to represent latent variables) that are related to variation in a response variable (Garthwaite 1994,Carrascal et al. 2009). Cross-validation can then be used to prevent overfitting of the data. We implemented this analysis using the PLS package in R (Mevik and Wehrens 2007). Cross-validation was used to select components that were related to response variables using the ‘leave-one-out’ method employed in the plsr() function. Twenty-three potential predictors were included in the PLSR analysis, including all of the WL variables described above, annual degree days for 0, 5, 10 and 15°C, annual precipitation, annual mercury deposition and annual sulfate deposition. We ran an individual PLSR with all 23 of these potential predictor variables for each of the sites with more than 12 years of data (6 sites, 1 in each lake). Root mean square error of prediction (RMSEP) was used to determine whether the inclusion of a component was strongly supported by the data: If inclusion of a component lowered the RMSEP, then the component was considered strongly supported and was included in the model (Mevik and Wehrens 2007). In cases where the RMSEP was at a minimum in the model with no components, we assumed no strong associations existed between the components and the response variable. Examples of the code are provided in the statistical appendix.

A hydrologic model by Thompson (2013)was previously developed that uses weather data as inputs and predicts what the WLs would be given different WL management scenarios. For the period 2000–2014, this hydrologic model can be used to estimate what the WLs would have been if the 1970 Rule Curve had been retained. The same model was also used to estimate what the WLs would have been if the 2000 Rule Curve were used. During the 2000–2014 period, the 2000 Rule Curve was used, but instead of using actual WL values, we used the model estimated WL values for this exercise for consistency. Thus, this hydrologic model provides estimated WL fluctuation estimates for a given WL management strategy. These WL estimates were then used as the predictor data in PLSR models that had strongly supported components to estimate young-of-year Yellow Perch mercury content. Code for making the predictions is included in the statistical appendix.

To incorporate potential error in the models, we used jackknife methods to make estimates of mercury in young-of-year Yellow Perch from 2000–2014. Essentially, the jackknife approach involved dropping one of the observations (observations here are the annual means at a site), refitting the PLSR model using the remaining observations, making predictions using that model, and repeating the process until estimates had been made from all the possible combinations of data that lacked one observation.

Correlations between individual WL variables and young-of-year Yellow Perch mercury content were performed using the Bayesian First Aid package (Bååth 2014) for each of the 13 sites sampled (both those sampled long-term and those sampled only during 2001–2003 and 2013–2015). Correlation coefficients were calculated with credible intervals. If those credible intervals do not overlap zero, then we consider it strong evidence that the correlation is non-zero (McCarthy 2007).

## Results

### Yellow Perch Hg content from separate sites within the same lake co-varies

Pearson’s correlation coefficients between sites within a single lake were large (all were >0.80) and only in Lake Kabetogama did the 95% credible interval of a Pearson’s. overlap zero (Figure 1). This indicates that within a particular lake, different sites tend to vary the same way over time, although these correlations are estimates from only 4–6 years (Figure 1). Most correlations between Yellow Perch Hg content and individual WL parameters were similar within a particular lake, with all 95% credible intervals overlapping from within a particular lake (Table 2). For example, the 95% credible interval of the Pearson’s. estimate for the association between Yellow Perch Hg content and Max WL in Little Vermilion Lake Site 1 (−0.42 to 0.68) and Site 2 (-0.90 to 0.71) had broad overlap (Table 2).

**Table 2.** Correlation coefficients (with 95% credible intervals) between characteristics of water level variation and whole young-of-year Yellow Perch total mercury content (per unit dry mass). Max WL-maximum annual WL, WLR-Annual water level rise, Amax WL-change in maximum WL since last year, NA-indicates the model would not converge. *-sites used in the partial least squared regression analysis. Bold indicates a non-zero correlation co-efficient.

| Lake | Site | Years of data | Max WL | WLR | Spring WLR | Δmax WL |
| --- | --- | --- | --- | --- | --- | --- |
| Crane Lake | 1 | 2001-03, 2013-15 | 0.56 (-0.30 to 0.95) | 0.50 (-0.36 to 0.94) | 0.53 (-0.35 to 0.94) | **0.80 (0.16 to 0.98)** |
| Crane Lake | 2* | 2001-10, 2013-15 | **0.57 (0.05 to 0.87)** | **0.54 (0.02 to 0.86)** | **0.57 (0.09 to 0.86)** | **0.57 (0.07 to 0.87)** |
| Lake Kabetogama | 1* | 2001-10, 2013-15 | 0.49 (-0.04 to 0.83) | 0.42 (-0.12 to 0.8) | 0.28 (-0.28 to 0.72) | 0.33 (-0.24 to 0.76) |
| Lake Kabetogama | 2 | 2001-03, 2013-15 | 0.67 (-0.11 to 0.97) | **0.75 (0.07 to 0.98)** | 0.63 (-0.17 to 0.96) | 0.66 (-0.15 to 0.97) |
| Little Vermilion Lake | 1* | 2001-03, 2013, 2015 | 0.17 (-0.42 to 0.68) | 0.23 (-0.35 to 0.71) | 0.35 (-0.23 to 0.78) | 0.35 (-0.25 to 0.78) |
| Little Vermilion Lake | 2 | 2001-03, 2013, 2015 | -0.21 (-0.90 to 0.71) | -0.20 (-0.92 to 0.70) | -0.17 (-0.88 to 0.73) | 0.12 (-0.75 to 0.86) |
| Namakan Lake | 1* | 2001-10, 2013-15 | **0.72 (0.33 to 0.92)** | **0.61 (0.14 to 0.88)** | **0.59 (0.11 to 0.87)** | **0.53 (0.02 to 0.85)** |
| Namakan Lake | 2 | 2001-03, 2013-15 | **0.80 (0.19 to 0.99)** | 0.72 (-0.06 to 0.98) | 0.69 (-0.10 to 0.97) | **0.88 (0.41 to 0.99)** |
| Rainy Lake | 1* | 2001-10, 2013-15 | **0.82 (0.51 to 0.96)** | **0.88 (0.66 to 0.97)** | **0.82 (0.52 to 0.95)** | **0.51 (0.01 to 0.84)** |
| Rainy Lake | 2 | 2001-03, 2015 | 0.56 (-0.56 to 0.98) | (NA) | 0.46 (-0.70 to 0.98) | 0.62 (-0.55 to 0.99) |
| Sand Point Lake | 0* | 1997, 1999, 2000-10, 2013-15 | **0.64 (0.22 to 0.88)** | **0.58 (0.14 to 0.86)** | **0.59 (0.17 to 0.85)** | **0.68 (0.31 to 0.90)** |
| Sand Point Lake | 1 | 2001-03, 2013-15 | 0.70 (-0.06 to 0.97) | 0.66 (-0.19 to 0.96) | 0.67 (-0.16 to 0.97) | **0.80 (0.18 to 0.99)** |
| Sand Point Lake | 2 | 2001-03, 2013-15 | 0.59 (-0.28 to 0.96) | 0.60 (-0.24 to 0.96) | 0.65 (-0.17 to 0.97) | **0.75 (0.07 to 0.98)** |

**Figure 1.**
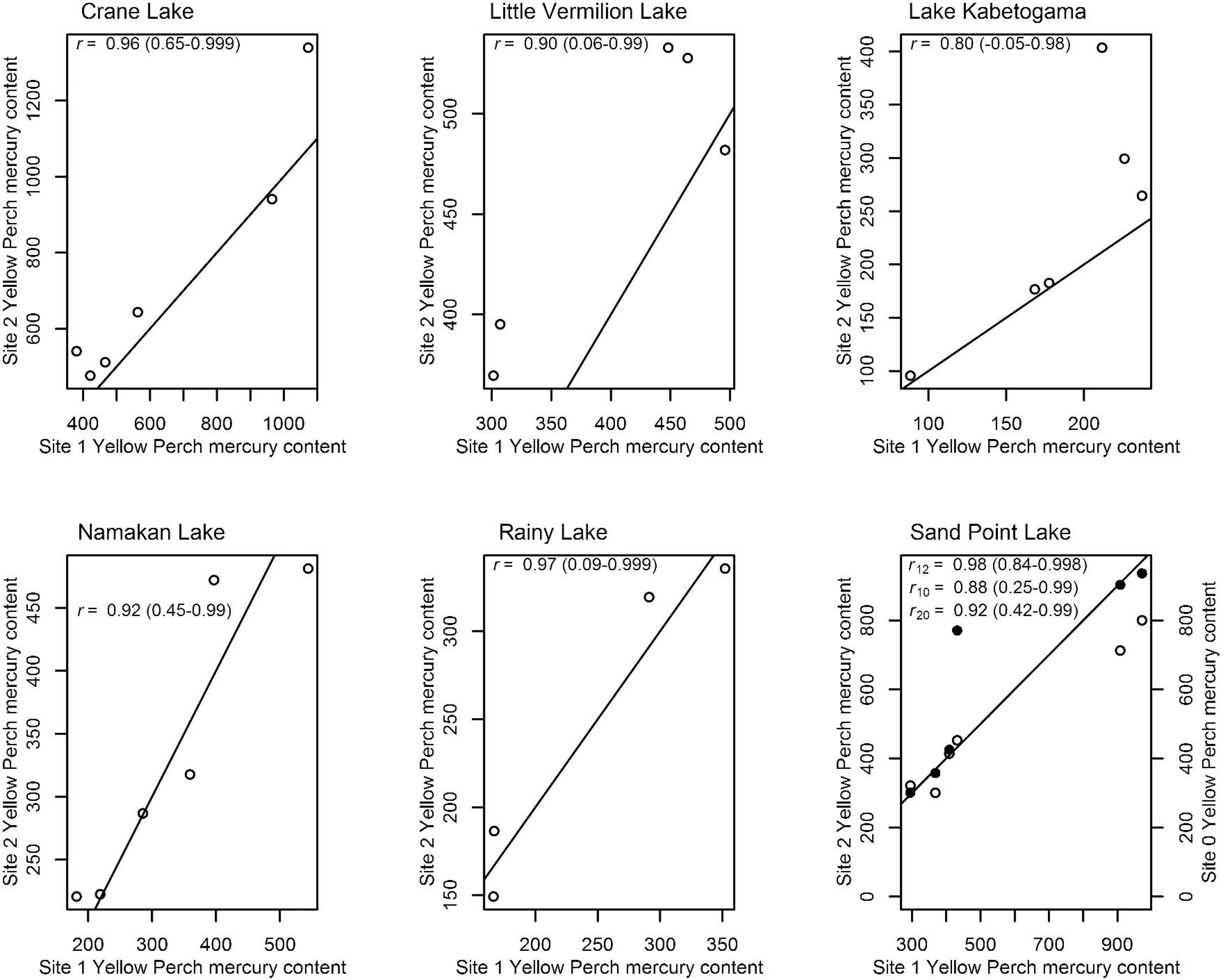
Annual variation in young-of-year whole fish Yellow Perch mean total mercury content in multiple sites from the same lake. Units for Yellow Perch mercury content are ng g^−1^ dry mass. The solid line is a 1:1 line. Pearson’s correlation coefficients (.) are reported with 95% confidence intervals. Each point is a separate year. For Sand Point Lake, closed circles are data from Site 0, open circles are from Site 2.

Bayesian estimates of correlation coefficients between annual variation in WL and young-of-year Yellow Perch Hg content suggests substantial spatial variation in the magnitude of WL effects. For example, the correlation between Yellow Perch Hg content and WLR was 0.88 (0.66 to 0.97) in the Rainy Lake site with the most data, but was indistinguishable from zero in the Little Vermilion Lake with the most data (0.23 [-0.35 to 0.71]; Table 2).

### Latent variables related to water level fluctuations strongly influence Yellow Perch Hg content

Partial least squared regression analysis suggested strong associations between WL fluctuations and young-of-year Yellow Perch mercury content in five of the six study lakes (Supplemental Tables 1–6, Figure 2). Only sites with 12+ years of data were used for this analysis. For each of these five lakes (Crane Lake, Lake Kabetogama, Namakan Lake, Rainy Lake and Sand Point Lake) the first component was very similar (Table 3). The first component explained between 42.9–79.2% of the variation in Yellow Perch mercury and always included the Spring WL rise, with greater rise leading to greater mercury content (Table 3). Only WL variables had correlations of >0.30 to this first component, and even then only a few crossed this threshold in each lake. Lake Kabetogama, Namakan Lake and Rainy Lake had second components that were supported by the cross-validation as well, and these second components were all positively associated with degree days (Table 4). None of the lakes had 3^rd^ components that were supported by cross-validation.

**Figure 2.**
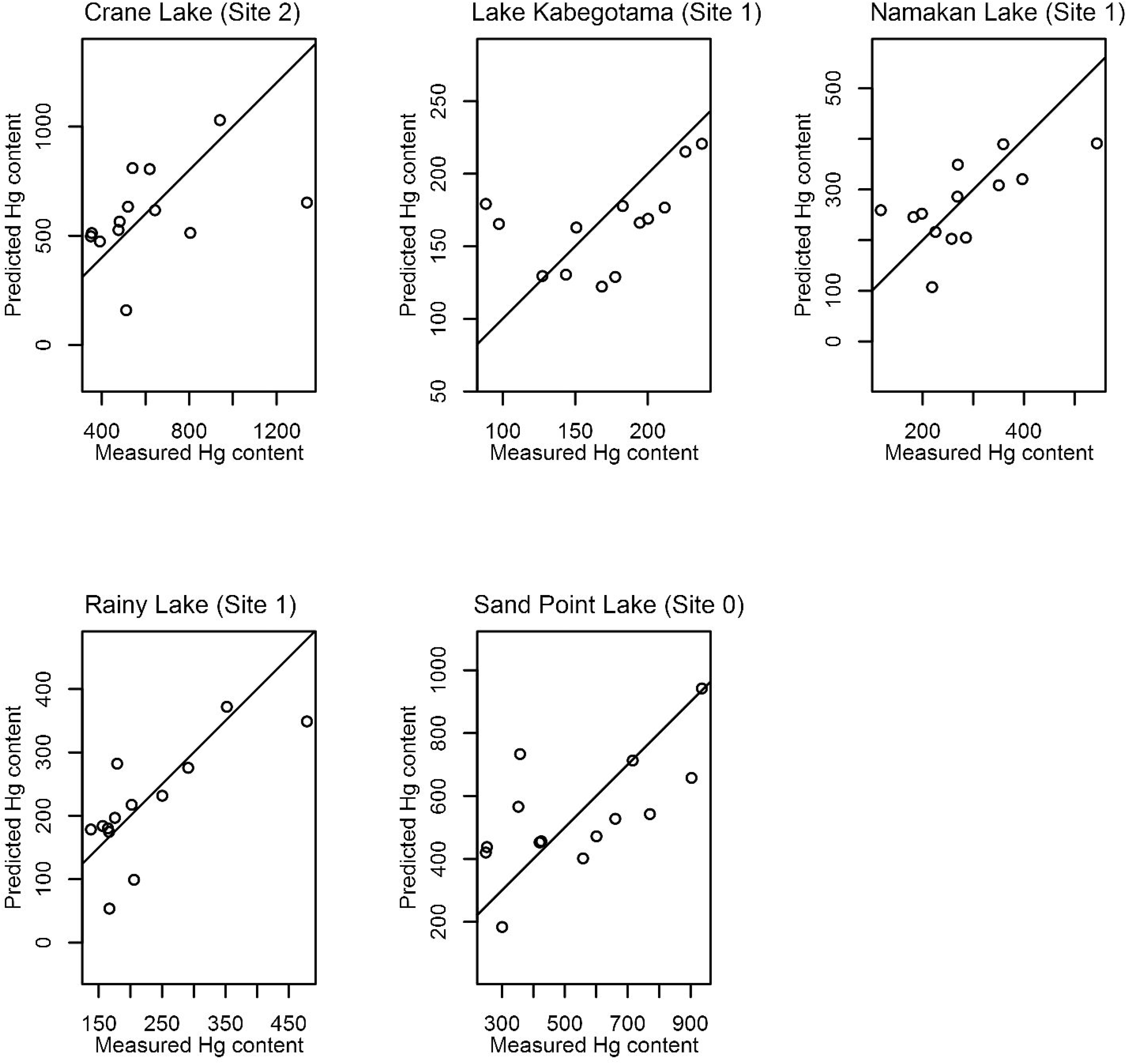
Mean measured and predicted whole fish young-of-year Yellow Perch total mercury (ng g^−1^ dry mass). Predictions are derived from partial least squared regression models that were validated using cross-validation. Crane Lake and Sand Point Lake models included 1 component, other lakes included 2 components (see Tables 3 and 4). The solid line is a 1:1 line.

**Table 3.** The first component of a partial least squared regression analysis relating water level (WL) fluctuations, degree days, precipitation, sulfate deposition and mercury deposition to total mercury content (per unit dry mass) in whole young-of-year Yellow Perch (collected at the end of September). Loadings are equivalent to a correlation between the component and the individual variable. Only loadings of 0.3 or more are shown. Cross-validation supported the inclusion of each of these components. Each of these sites had >12 years of data. Spring refers to April-June; Summer refers to July-September. No components were supported for Little Vermilion Lake.

|  | Crane Lake<br>(Site 2) | Lake<br>Kabetogama<br>(Site 1) | Namakan Lake<br>(Site 1) | Rainy Lake<br>(Site 1) | Sand Point<br>Lake (Site 0) |
| --- | --- | --- | --- | --- | --- |
| % Variance Explained<br>(predictors) | 46.5 | 42.8 | 46.4 | 48.1 | 47.6 |
| % Variance Explained (mean<br>Yellow Perch Hg ng per g DW) | 54.2 | 42.9 | 61.7 | 79.2 | 62.3 |
| *Loadings* |  |  |  |  |  |
| Annual maximum WL | 0.30 | 0.35 | 0.31 | 0.30 | - |
| Annual WL rise | - | 0.34 | 0.31 | - | 0.30 |
| Change in maximum WL from<br>last year | - | - | - | - | - |
| Annual minimum WL | - | - | - | - | - |
| Annual mean WL | - | - | - | - | - |
| Annual standard deviation WL | - | - | - | - | - |
| Spring maximum WL | 0.30 | 0.32 | 0.31 | 0.30 | - |
| Spring WL rise | 0.30 | 0.31 | 0.30 | 0.30 | 0.30 |
| Spring minimum WL | - | - | - | - | - |
| Spring mean WL | - | - | - | - | - |
| Spring standard deviation WL | - | - | - | - | - |
| Summer maximum WL | 0.30 | 0.33 | - | 0.30 | - |
| Summer WL rise | - | 0.32 | - | - | - |
| Summer minimum WL | - | - | - | - | - |
| Summer mean WL | - | 0.33 | - | - | - |
| Summer standard deviation WL | - | 0.33 | - | - | - |
| Degree days (0°C) | - | - | - | - | - |
| Degree days (5°C) | - | - | - | - | - |
| Degree days (10°C) | - | - | - | - | - |
| Degree days (15°C) | - | - | - | - | - |
| Precipitation (mm) | - | - | - | - | - |
| Sulfate deposition | - | - | - | - | - |
| Mercury deposition | - | - | - | - | - |

**Table 4.** The second component of a partial least squared regression analysis relating water level (WL) fluctuations, degree days, precipitation, sulfate deposition and mercury deposition to total mercury content (per unit dry mass) in whole young-of-year Yellow Perch (collected at the end of September). Loadings are equivalent to a correlation between the component and the individual variable. Only loadings of 0.3 or more are shown. Cross-validation supported the inclusion of each of these components. Each of these sites had >12 years of data. Spring refers to April-June; Summer refers to July-September. Crane Lake and Sand Point Lake did not have strongly supported second components, while Little Vermilion Lake had no strongly supported components.

|  | Lake<br>Kabetogama<br>(Site 1) | Namakan Lake<br>(Site 1) | Rainy Lake<br>(Site 1) |
| --- | --- | --- | --- |
| % Variance Explained<br>(predictors) | 19.4 | 14.2 | 21.2 |
| % Variance Explained (mean<br>Yellow Perch Hg ng per g DW) | 23.6 | 21.4 | 8.0 |
| Annual maximum WL | - | - | - |
| Annual WL rise | - | - | - |
| Change in maximum WL from<br>last year | - | - | - |
| Annual minimum WL | - | 0.38 | -0.34 |
| Annual mean WL | - | - | - |
| Annual standard deviation WL | - | - | - |
| Spring maximum WL | - | - | - |
| Spring WL rise | - | - | - |
| Spring minimum WL | - | 0.36 | -0.34 |
| Spring mean WL | - | - | - |
| Spring standard deviation WL | -0.31 | - | - |
| Summer maximum WL | - | - | - |
| Summer WL rise | - | - | - |
| Summer minimum WL | - | - | -0.31 |
| Summer mean WL | - | - | - |
| Summer standard deviation | - | - | - |
| Degree days (0°C) | 0.36 | 0.34 | - |
| Degree days (5°C) | 0.38 | 0.33 | 0.31 |
| Degree days (10°C) | 0.40 | 0.35 | 0.32 |
| Degree days (15°C) | 0.41 | 0.38 | 0.30 |
| Precipitation (mm) | - | - | - |
| Sulfate deposition | - | - | - |
| Mercury deposition | - | - | - |

### Statistical models suggest water level management changes would cause changes in Yellow Perch Hg content

The five strongly supported PLSR models were used to model the effects of the two WL management scenarios. These were the models previously described in Tables 2 and 3, and included a 1 component model for Crane Lake and Sand Point Lake, and 2-component models for Lake Kabetogama, Namakan Lake and Rainy Lake (as mentioned above, no components were included in models for Little Vermilion Lake). The two WL management scenarios included one scenario that used the 1970 Rule Curve and another that used the 2000 Rule Curve prescribed by the International Joint Commission for these lakes (IJC 2000). Annual variation in young-of-year Yellow Perch mercury content was similar under either management scenario; however, predictions for Crane Lake, Namakan Lake, Sand Point Lake and Lake Kabetogama tended to be higher under the 1970 Rule Curve scenario (Figure 3). For Rainy Lake, both management scenarios yield similar annual variation in fish mercury content (Figure 3), which is not surprising since the 1970 and 2000 Rule Curves for Rainy Lake are almost identical (IJC 2000).

**Figure 3.**
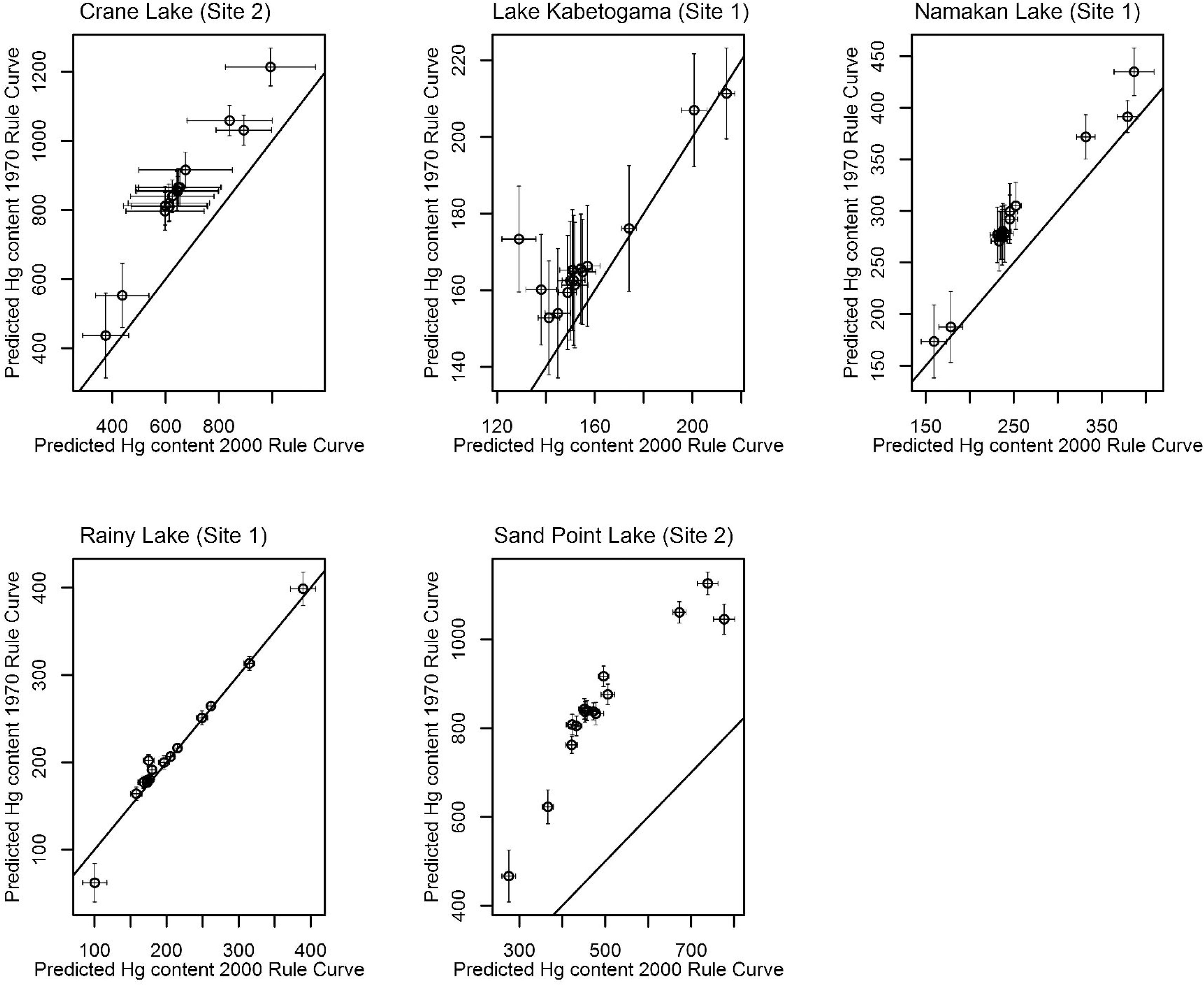
Predicted whole fish young-of-year Yellow Perch total mercury content (ng g^−1^ dry mass) in response to changes in water level (WL) management. Predictions are based on partial least squared regression (PLSR) models and two WL management scenarios generated from a hydrological model spanning 2000–2014. Points are means of jackknifed PLSR models (error bars are standard deviations from the jackknifed estimates).

## Discussion

In previous studies on the lakes in this analysis, year-to-year changes in maximum WL appeared to be the biggest driver of annual variation in young-of-year Yellow Perch mercury content (Sorensen et al. 2005,Larson et al. 2014). Year-to-year change in maximum WL was strongly associated with annual variation in Yellow Perch mercury content in this analysis as well (as indicated by correlation coefficients), but other WL metrics were equally important. However, in some lakes these associations were absent (Little Vermilion Lake) and in some the associations were very strong (Rainy Lake).

Use of partial least squared regression (PLSR) allowed us to incorporate many different WL metrics (and other environmental variables) into a more inclusive model than had been used in previous analyses (Larson et al. 2014). The PLSR models demonstrated some lake-specific difference analogous to those seen in previous studies. For example, PLSR could not identify any WL-influenced latent variables that might be driving annual variation in Little Vermilion Lake, consistent with Larson et al.’s (2014) finding that WL-fish mercury associations in that lake were weak. But PLSR also was able to identify some commonalities among lakes that the previous study did not detect. For example, Yellow Perch mercury content in Lake Kabetogama appeared unrelated to WL effects in Larson et al.’s (2014) analysis, but in the PLSR analysis a latent component correlated to a few WL metrics appeared to be strongly associated with annual variation in fish mercury content. A very similar latent component appeared in every other lake (except Little Vermilion Lake). Although it is still unclear why quantitative differences exist in the magnitude of these WL-fish mercury associations among lakes, qualitatively these associations appear more similar among lakes than they did in the less inclusive modeling approach (Larson et al. 2014). These spatial differences also appear to be driven by lake-wide differences, since different sites within the same lake appeared to behave similarly.

Overall the PLSR models suggested that WL rise (in Spring and in some lakes annual) was positively associated with young-of-year Yellow Perch mercury content. These are WL characteristics that were altered in five of these lakes (all but Rainy Lake) by the 2000 Rule Curves prescribed by the International Joint Commission. (IJC 2000). These rule curves reduced the winter drawdown in Namakan Reservoir by approximately 1 m, whereas changes in Rainy Lake were minimal. When comparing the modeled WLs in these lakes using both the new and old regulations, differences emerge in the estimated fish mercury content. For Crane Lake and Sand Point Lake, the models clearly suggest that fish mercury content would have been higher from 2000–2014 if the 1970 Rule Curve had been used. For other lakes the models suggest no change (Rainy Lake) or suggest slight increases (Lake Kabetogama and Namakan Lake) from 2000–2014 if the 1970 Rule Curve had been in place.

Methylmercury contamination of fisheries is a major management concern (Sandheinrich and Wiener 2011), and it appears that certain water level management strategies could possibly influence the magnitude of this problem. Previous research has shown that differences in reservoir construction, initial inundation, and overall purpose may influence mercury content in fish (Dembkowski et al. 2014b,Willacker et al. 2016), but here we show that different water level management regimes in established impounded lakes could be associated with changes in fish mercury content. Management for methylmercury contamination is rarely the only consideration in WL management, but models such as those developed in this study could be used to evaluate different WL management scenarios so that consideration for multiple uses of a particular waterway could be incorporated.

## Acknowledgments

We would like to thank two U.S. Geological Survey reviewers for their helpful comments on this manuscript. Any use of trade, product, or firm names is for descriptive purposes only and does not imply endorsement by the U.S. Government.

## Literature Cited.

Bååth, R. 2014. Bayesian First Aid. https://github.com/rasmusab/bayesian_first_aid.

Carrascal, L. M., I. Galván, and O. Gordo. 2009. Partial least squares regression as an alternative to current regression methods used in ecology. Oikos 118:681–690.

Chezik, K. A., N. P. Lester, P. A. Venturelli, and K. Tierney. 2014. Fish growth and degree-days I: selecting a base temperature for a within-population study. Canadian Journal of Fisheries and Aquatic Sciences 71:47–55.

Christensen, V. G., J. H. Larson, R. P. Maki, M. B. Sandheinrich, M. E. Brigham, C. Kissane, and J. F. LeDuc. 2017. Lake levels and water quality in comparison to fish mercury body burdens, Voyageurs National Park, Minnesota, 2013–15. Scientific Investigations Report 2016–5175. Reston, Virginia.

Dembkowski, D. J., S. R. Chipps, and B. G. Blackwell. 2014a. Response of walleye and yellow perch to water-level fluctuations in glacial lakes. Fisheries Management and Ecology 21:89–95.

Dembkowski, D. J., S. R. Chipps, and B. G. Blackwell. 2014b. Response of walleye and yellow perch to water-level fluctuations in glacial lakes. Fisheries Management and Ecology 21:89–95.

Driscoll, C. T., Y.-J. Han, C. Y. Chen, D. C. Evers, K. F. Lambert, T. M. Holsen, N. C. Kamman, and R. K. Munson. 2007. Mercury Contamination in Forest and Freshwater Ecosystems in the Northeastern United States. BioScience 57:17–28.

Garthwaite, P. 1994. An interpretation of partial least squares. Journal of the American Statistical Association 89:122–127.

Holmlund, C. M., and M. Hammer. 1999. Ecosystem services generated by fish populations. Ecological Economics 29:253–268.

IJC, (International Joint Commission). 2000. Supplementary Order to the Order Prescribing Method of Regulating the Levels of Boundary Waters. Pages 1–6.

Larson, J. H., R. P. Maki, B. C. Knights, and B. R. Gray. 2014. Can mercury in fish be reduced by water level management? Evaluating the effects of water level fluctuation on mercury accumulation in yellow perch (Perca flavescens). Ecotoxicology 23:1555–1563.

Mailman, M., L. Stepnuk, N. Cicek, and R. Bodaly. 2006. Strategies to lower methyl mercury concentrations in hydroelectric reservoirs and lakes: A review. The Science of the Total Environment 368:224–35.

Manly, B. F. J. 2005. Multivariate Statistical Analysis: A Primer. 3rd edition. CRC Press LLC, Boca Raton, FL.

McCarthy, M. 2007. Bayesian methods for ecology. Cambridge University Press, New York, New York, USA.

Mevik, B., and R. Wehrens. 2007. The pls package: principal component and partial least squares regression in R. Journal of Statistical Software 18.

Munthe, J., R. A. D. Bodaly, B. A. Branfireun, C. T. Driscoll, C. C. Gilmour, R. Harris, M. Horvat, M. Lucotte, and O. Malm. 2007. Recovery of Mercury-Contaminated Fisheries. Ambio 1:33–44.

NADP. 2012a. NADP Site Information (MN18).

http://nadp.sws.uiuc.edu/sites/siteinfo.asp?id=MN18&net=NTN.

NADP. 2012b. NADP Site Information (MN32).

http://nadp.sws.uiuc.edu/sites/siteinfo.asp?net=NTN&id=MN32.

R Development Core Team. 2014. R: A language and environment for statistical computing. R Foundation for Statistical Computing, Vienna, Austria.

Sandheinrich, M. B., and J. G. Wiener. 2011. Methylmercury in freshwater fish: Recent advances in assessing toxicity of environmentally relevant exposures. Pages 169–192 in W. N. Beyer and J. P. Meador, editors. Environmental Contaminants in Biota: Interpreting Tissue Concentrations. 2nd edition. CRC Press, Boca Raton, FL.

Scheuhammer, A. M., M. W. Meyer, M. B. Sandheinrich, and M. W. Murray. 2007. Effects of environmental methylmercury on the health of wild birds, mammals, and fish. Ambio 36:12–18.

Selin, N. E. 2009. Global Biogeochemical Cycling of Mercury: A Review. Annual Review of Environment and Resources 34:43–63.

Sorensen, J. A., L. W. Kallemeyn, and M. Sydor. 2005. Relationship between Mercury Accumulation in Young-of-the-Year Yellow Perch and Water-Level Fluctuations. Environmental Science & Technology 39:9237–9243.

Thompson, A. F. 2013. Rainy and Namakan Hydrologic Response Model Documentation.

Wiener, J. G., B. C. Knights, M. B. Sandheinrich, J. D. Jeremiason, M. E. Brigham, D. R. Engstrom, L. G. Woodruff, W. F. Cannon, and S. J. Balogh. 2006. Mercury in Soils, Lakes, and Fish in Voyageurs National Park ( Minnesota ): Importance of Atmospheric Deposition and Ecosystem Factors. Environmental Science & Technology 40:6261–6268.

Willacker, J. J., C. A. Eagles-Smith, M. A. Lutz, M. T. Tate, J. M. Lepak, and J. T. Ackerman. 2016. Reservoirs and water management influence fish mercury concentrations in the western United States and Canada. The Science of the Total Environment 568:739–748.

